# Characterization Of Hydrocarbon Utilizing Bacteria In Waste Engine Oil-Impacted Sites

**DOI:** 10.1101/2020.03.21.998872

**Authors:** A. M. Jones, I. I. James, P. S. Akpan, I.I. Eka, A. E. Oruk, A. A. Ibuot

## Abstract

The present work was aimed at isolating and identifying hydrocarbon utilizing bacteria from waste engine oil polluted soil, and assessing their hydrocarbon-utilizing ability. Materials and method: Eight bacterial species; *Corynebacterium kutscheri, Pseudomonas aeruginosa, Flavobacterium aquatile, Serratia odorifera, Micrococcus agilis, Staphylococcus aureus, Micrococcus luteus* and *Bacillus substilis* were isolated by selective enrichment technique and screened for hydrocarbon utilization capability in mineral salt media with waste engine oil as a sole carbon and energy source. Bacterial species showed varied hydrocarbon utilization during the 15 days of incubation. All isolates also showed variable emulsification ability. These results demonstrate the presence of indigenous bacteria in hydrocarbon polluted soils and the potential towards remediation of hydrocarbons.

## 1. INTRODUCTION

Petroleum utilization as fuels and petroleum products leads to severe environmental pollution [1]. Large-scale accidental spills pose a great threat to the ecosystem [2]. Soil pollution by petroleum hydrocarbons has been shown to produce pronounced changes in the physicochemical and microstructure of the oil-contaminated soil [3]. This affects parameters such as soil porosity, bulk density, adsorbtion, etc [4, 5]. Fresh spills and/or high levels of pollutants may often result in the reduction of large sectors of soil microbial population, although soils with lower levels or old pollution may show increase in numbers and diversity of microorganisms [6,7].

The diversity and the number of microorganisms at a polluted soil site may assist in the characterization of such a site; such as toxicity of petroleum hydrocarbons to the microbiome, age of the spill and concentration of the pollutant [8]. Additionally, microorganism in soils exposed to hydrocarbon pollution usually exhibits a higher potential for biodegradation of such pollutant compounds than others with no history of such exposure. Percentages of hydrocarbonoclastic microbes are quite low in soil when there is no oil spill, but may increase 1,000 fold after oil spill [9].

Conventional remediation methods do not seem to be able to address this problem, or tends to aggravate the problem [10]. Mechanical methods such as incineration, excavation and/or burial in secure land fill, as well as a host of other chemical decomposition methods are expensive, time consuming and only relocates the pollution [11]. An efficient way of remediating the oil-contaminated sites could be employment of microorganisms, such as bacteria, microscopic algae, and fungi, isolated from polluted environments or enhanced from the organisms already present in the same environment [12, 13]. Waste engine oil-polluted soils also serve as a source of indigenous bacteria capable of hydrocarbon degradation.

The employment of microorganisms in the biodegradation of hydrocarbons over chemical or conventional treatment is preferred for many reasons; end products that are comparatively safer and cost-effectiveness [11]. This present study considered the indigenous bacterial communities in waste engine oil-polluted soil capable of utilizing waste engine oil.

## 2. MATERIALS AND METHODS

### Sample collection

Waste engine oil –contaminated soil samples used in this study were collected from three auto-mechanic workshops within the mechanic village, Uyo, Akwa Ibom State. Composite soil samples were obtained at each sampling point using a soil auger from 0-10 cm below the soil surface. The soils were labeled “Unpolluted” for the unpolluted sample, “MA” for the mechanic workshop 1 sample, “MB” for the mechanic workshop 2 sample, and “MC” for the mechanic workshop 3 sample. This was followed by bulking and transportation to the laboratory in sterile polythene bags within six hours for isolation of organisms.

### Physico-chemical analysis of soil samples

Some physicochemical parameters of the soil sample were analyzed as follows:

Soil pH was measured using HANNA Instruments Model 209 pH meter [14]. Moisture content was calculated on the basis of the air dry weight as described by AOAC [15]. Total organic carbon was calculated by weighing exactly 0.5 g of the soil sample into a flask, and 10 ml of 1.0 M K_2_Cr_2_O_7_ was added and swirled to mix. 20 ml conc. H_2_SO_4_ was added, swirled gently for a minute and allowed to stand for 20 minutes. The suspension was diluted to about 100 ml of the distilled water. 5 drops of o-phenanthroline indicator was added to each sample and was titrated with 0.5 M ferrous ammonium sulphate to a light blue end point. The reagent blank was also run and the titre values recorded, and used to calculate the organic carbon content [15].

Total hydrocarbon content was determined by first extracting hydrocarbons by acidifying 2 g of representative soil samples using H_2_SO_4_, and extracting upon addition of 20 ml of toluene in a seperatory funnel. The contents of the funnel were shaken, and allowed to settle into two layers. The absorbance of the supernatant (extract) was read at 420 nm with UNICAM UV/VIS spectrophotometer (Spectronic 20D). Readings were recorded from the spectrophotometer and using the determined curve to get the figure [16].

Phosphorus was determined using the ascorbic acid method as described by AOAC, [15] 50 ml of the soil dilution was pipetted into 250 ml Erlenmeyer flask, and 1 drop of phenolphthalein indicator added. Exactly 5 N H_2_SO_4_ (148 ml conc. H_2_SO_4_ in 100ml H_2_0) is added drop-wise to develop a red colour. Exactly 8 ml of combined reagents made up of 50 ml of 5 N H_2_SO_4_, 5 ml potassium antimonyl tartrate solution (1.372g potassium antimonyl tartrate in 500 ml distilled water); 15 ml ammonium molybdate solution (20g ammonium molybdate crystal in 500 ml distilled water); were added and thoroughly mixed, and allowed to stand for 20 minutes. The phosphorus content was determined by measuring the absorbance of the sample at 880 nm.

Nitrogen concentration was determined according to the methods of Bremmer and Mulvaney, [17]. One milliliter of the soil sample was introduced into standard kjeldahl flask containing 1.5 g CuSO_4_, and 1.5 g Na_2_SO_4_ as catalyst, alongside concentrated H_2_SO_4_. The flask was gently heated on a heating mantle, taking care to prevent frothing. The solution was transferred after heating to a 100 ml standard flask and made up to the mark with distilled water. A portion of this digest was pipetted into a semi micro-kjeldahl distillation apparatus and treated with 30 ml 0of 40 % NaOH solution. The ammonia evolved was steam-distilled into a 100 ml conical flask containing 10 ml solution of saturated boric acid to which 4 drops of Tashirus indicator had been previously added. The tip of a condenser was immersed into the boric acid solution and distillation continued till about two-thirds of the original volume was obtained. The tip of the condenser was finally rinsed with a few milliliters of distilled water. The distillate was then titrated with 0.1N HCl until a purple-pink end point was observed. A blank determination was also carried out in a similar manner without the sample, and calculation done as follows; 

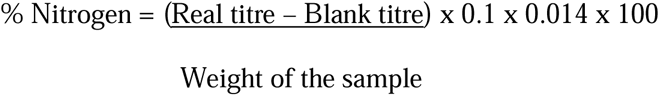

### Enumeration of total heterotrophic bacteria

Total heterotrophic bacterial population in the soil samples was enumerated by adopting the standard plate counts technique using spread plate method as described by Ogunbayo *et al.*, [18]. These involved spreading aliquots of a serially diluted 0.1 ml of 10^−5^ dilutions of the soil sample suspension on nutrient agar plates and the plates were incubated at 30 °C for 24 hours. Similar aliquots were also incubated in minimal salt agar plates containing used engine oil as the sole source of carbon and energy. The plates were all incubated aerobically at 30 °C. The percentages of hydrocarbon-utilizing bacteria (HUB) relative to the total heterotrophic counts were noted.

### Enumeration of hydrocarbon utilizing bacteria

Oil-utilizing bacteria were isolated from the polluted soil samples by enrichment in mineral salt medium (MSM) modified from Okpokwasili and Nwosu, [19], using waste engine oil as sole carbon source [19]. The soil samples were sieved using a 2mm mesh sieve. 10 g of the sieved soil samples was inoculated into 100ml sterile MSM. 1 ml of the waste engine oil was added to the medium as a sole source of carbon and energy, and the culture incubated on a rotary shaker at 170 rpm for 1 week.

The enrichment procedure was repeated for three cycles. At the end of each enrichment cycle, 1ml of the culture was diluted serially 10-fold down the gradient to 10^−5^ and plated. Pure cultures of the isolates were obtained by plating 1ml of the 10^−5^ dilution of the third enrichment cycle onto MSM agar plates, and incubating at 30 °C (± 2) for 48 hours. Pure cultures obtained by this procedure were stored in slants at 4°C until further identification.

### Characterization and identification of bacteria

Isolates were identified based on colonial characteristics, Gram’s reaction and cell biochemical reactions as described by Cheesbrough [20] Identification used the taxonomic schemes of Holt *et al*. [21].

### Hydrocarbon utilization screening of bacteria

To determine the ability of the isolates to utilize engine oil as sole source of carbon and energy, growth patterns of isolates in mineral salt medium in the presence of 1% (v/v) of the waste engine oil (5.0 mL in 100 mL MSM) was determined according to Onuoha *et al.*[22]. Waste engine oil-augmented MSM was dispensed into 250 ml Erlenmeyer, and inoculated with 0.1ml of 24h cultures of the bacterial isolates. Incubation was done at 30 °C for 15 days. Growth patterns were determined monitoring changes in pH, optical density and total viable count at 5-day intervals during the incubation. The pH of the medium was measured using the pH meter (HANNA Instruments). Growth was also monitored by measuring the optical density (OD) at 600 nm using the spectrophotometer (Spectrumlab). Total viable counts of the cultures were obtained by incubation of 0.1 ml of the cultures using spread plate technique on nutrient agar plates at 30 °C for 24 hours.

### Emulsification activity of bacteria

The emulsification index (E_24_) of the isolates was determined according to the methods of Ganesh and Lin, [23], by adding 1ml of waste engine oil to the same amount of culture media as used for degradation assay, mixing the vortex for 2 minutes and leaving to stand for 24 h. The percentage of emulsification index was obtained as follows; 

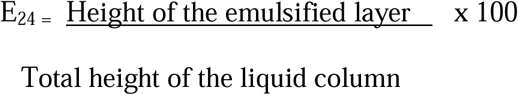

## 3. RESULTS

### Physicochemical properties of soil samples

The results of the physicochemical analysis of the different soil sample are shown in Table 1. The high amounts of organic carbon (5.32 ± 2.65 % in MA, 9.79 ± 0.51 % in MB and 7.29 ± 3.09 % in MC), and THC (2933.76 ± 404..27 mg/kg in MA, 3122.72 ± 131.00 mg/kg in MB and 3202.61± 675.07 mg/kg in MC), compared to the unpolluted soil sample (3.7 ± 2.43 % organic carbon content and 39.97± 13.49 mg/kg THC) is indicative of heavy pollution of the mechanical workshop samples with petroleum hydrocarbons.

Soil samples from mechanical workshop 1 contained higher amounts of nitrates (0.25 ± 0.03 mg/g), while samples from mechanical workshop 2 contained the highest amounts of phosphates (10.74 ± 0.88 mg/g) and THC (3202.61± 675.07 mg/kg). pH values of the soil samples indicates all soil samples as moderately acidic to acidic (5.78 to 6.79).

**Table 1:** Physicochemical properties of different soil samples contaminated with waste engine oil.

| Parameters | Unpolluted | Polluted |  |  |
| --- | --- | --- | --- | --- |
|  | (control) | MA | MB | MC |
| pH | 6.79 ± 0.24 | 6.18 ± 0.13 | 5.78 ± 0.03 | 6.44 ± 0.07 |
| Nitrate (mg/g) | 0.14 ± 0.02 | 0.25 ± 0.03 | 0.23 ± 0.02 | 0.20 ± 0.02 |
| Phosphate (mg/g) | 6.65 ± 1.72 | 6.92 ± 2.92 | 10.74 ± 0.88 | 8.49 ± 1.9 |
| Moisture content (%) | 32.25 ± 9.30 | 43.09 ± 0.65 | 45.13 ± 7.02 | 27.72 ± 9.09 |
| Organic carbon content (%) | 3.7 ± 2.43 | 5.32 ± 2.65 | 9.79 ± 0.51 | 7.29 ± 3.09 |
| THC (mg/kg) | 39.97 ± 13.49 | 2933.76 ± 404.27 | 3202.61 ± 675.07 | 3122.72 ± 131.00 |

Values represent means of triplicate determinations ± SD.
MA = Mechanic Workshop 1, MB = Mechanic Workshop 2, MC = Mechanic Workshop 3
THC = Total hydrocarbon content

### Bacterial count of soil samples

The total heterotrophic bacterial count and hydrocarbon utilizing bacterial count of the original soil samples is shown in Table 2. A higher THB count was recorded in polluted soil samples (4.4 ± 1.90 × 10^7^ cfu/g from MA sample, 6.0 ± 0.23 ×10^7^ cfu/g from MB sample and 4.5 ± 0.03 ×10^7^ cfu/g from MC sample) than in the unpolluted soil sample (1.9 ×10^7^ cfu/g).

**Table 2:**
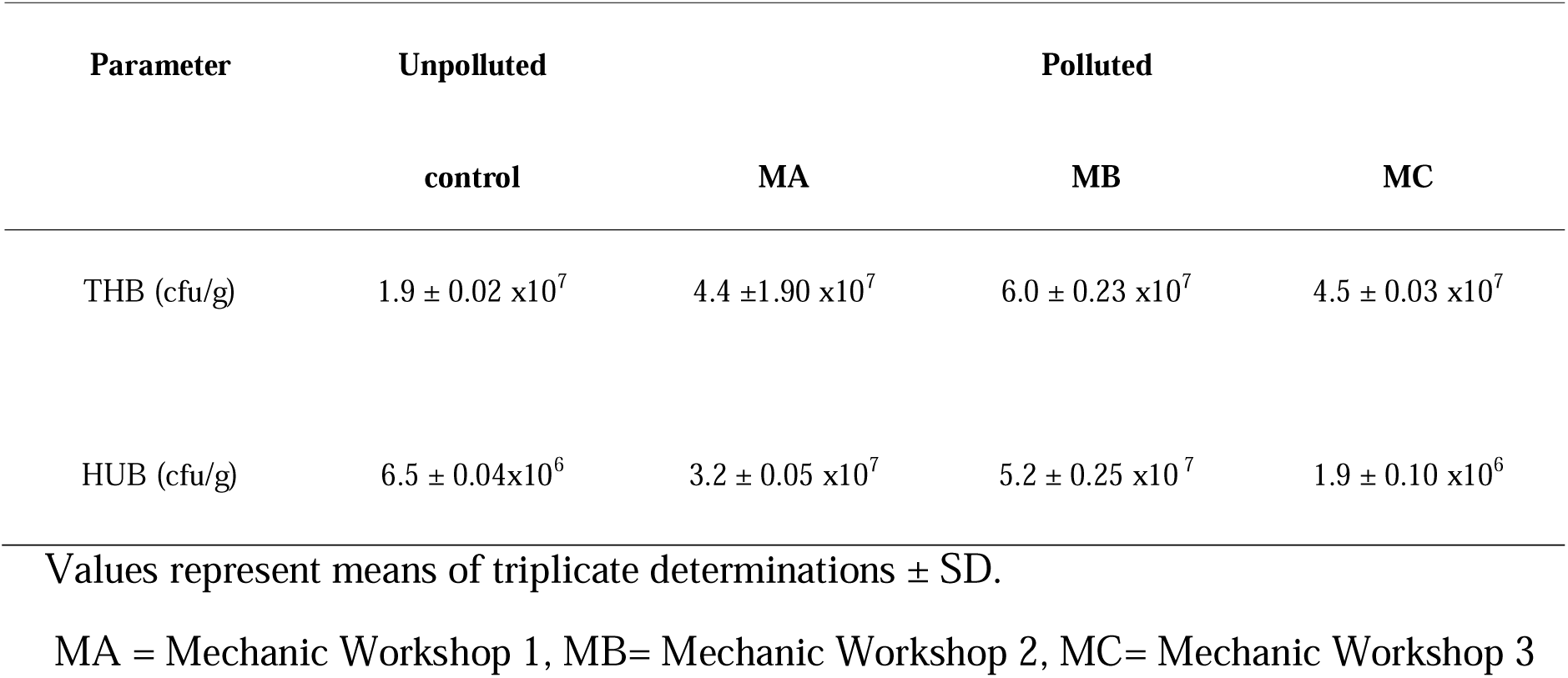
Bacterial count of soil samples contaminated with waste engine oil.

| Parameter | Unpolluted | Polluted |  |  |
| --- | --- | --- | --- | --- |
|  | control | MA | MB | MC |
| THB (cfu/g) | 1.9 ± 0.02 x10<sup>7</sup> | 4.4 ± 1.90 x10<sup>7</sup> | 6.0 ± 0.23 x10<sup>7</sup> | 4.5 ± 0.03 x10<sup>7</sup> |
| HUB (cfu/g) | 6.5 ± 0.04x10<sup>6</sup> | 3.2 ± 0.05 x10<sup>7</sup> | 5.2 ± 0.25 x10<sup>7</sup> | 1.9 ± 0.10 x10<sup>6</sup> |

Values represent means of triplicate determinations ± SD.
MA = Mechanic Workshop 1, MB= Mechanic Workshop 2, MC= Mechanic Workshop 3

Higher THB (6.0 ± 0.23 ×10^7^ cfu/g) and HUB (5.2 ± 0.25 ×10^7^ cfu/g) counts were observed in the MB sample than in other similar polluted samples indicative of its extent of pollution. Hydrocarbon utilising bacterial counts were slightly lower in all samples than the corresponding heterotrophic bacterial counts.

### Characterization and identification of bacteria

Identified bacterial isolates were *Corynebacterium kutscheri, Pseudomonas aeruginosa, Micrococcus agilis, Flavobacterium aquatile, Staphylococcus aureus, Micrococcus luteus, Serratia odorifera* and *Bacillus substilis* as shown on Table 3 and 4.

**Table 3:** Biochemical characteristics of bacterial isolates from soil contaminated with waste engine oil.

| Isolates | Isolate code | Spore | Catalase | Motility | Oxidase | Litmus reaction | Urease | Gelatin | Citrate | Glucose | Maltose | Manitol | Lactose | Sucrose | Arabinose | Xylose |
| --- | --- | --- | --- | --- | --- | --- | --- | --- | --- | --- | --- | --- | --- | --- | --- | --- |
| 1 | MA3, MB1, MC4 | - | + | - | - | - | - | - | + | A | A | A | - | - | - | AG |
| 2 | MA2, MB2 | - | + | + | + | + | - | + | - | - | A | A | A | - | A | A |
| 3 | MA1 | - | + | - | - | - | - | - | - | A | - | A | - | - | AG | AG |
| 4 | MA4 | - | + | - | + | + | - | - | - | A | A | - | A | A | - | - |
| 5 | MB3, MC1, MC2, MA5 | - | + | - | - | + | - | - | - | - | - | - | - | A | - | - |
| 6 | MA6 | - | + | - | + | - | - | - | - | - | - | A | - | A | - | - |
| 7 | MC3 | - | + | + | - | - | + | + | + | A | A | A | A | A | A | - |
| 8 | MB4, MC5 | + | + | + | - | + | - | + | + | A | A | A | - | A | A | AG |

**Keys:** AG = acid and gas production; A = acid production only; + = positive reaction; - = negativ

**Table 4:**
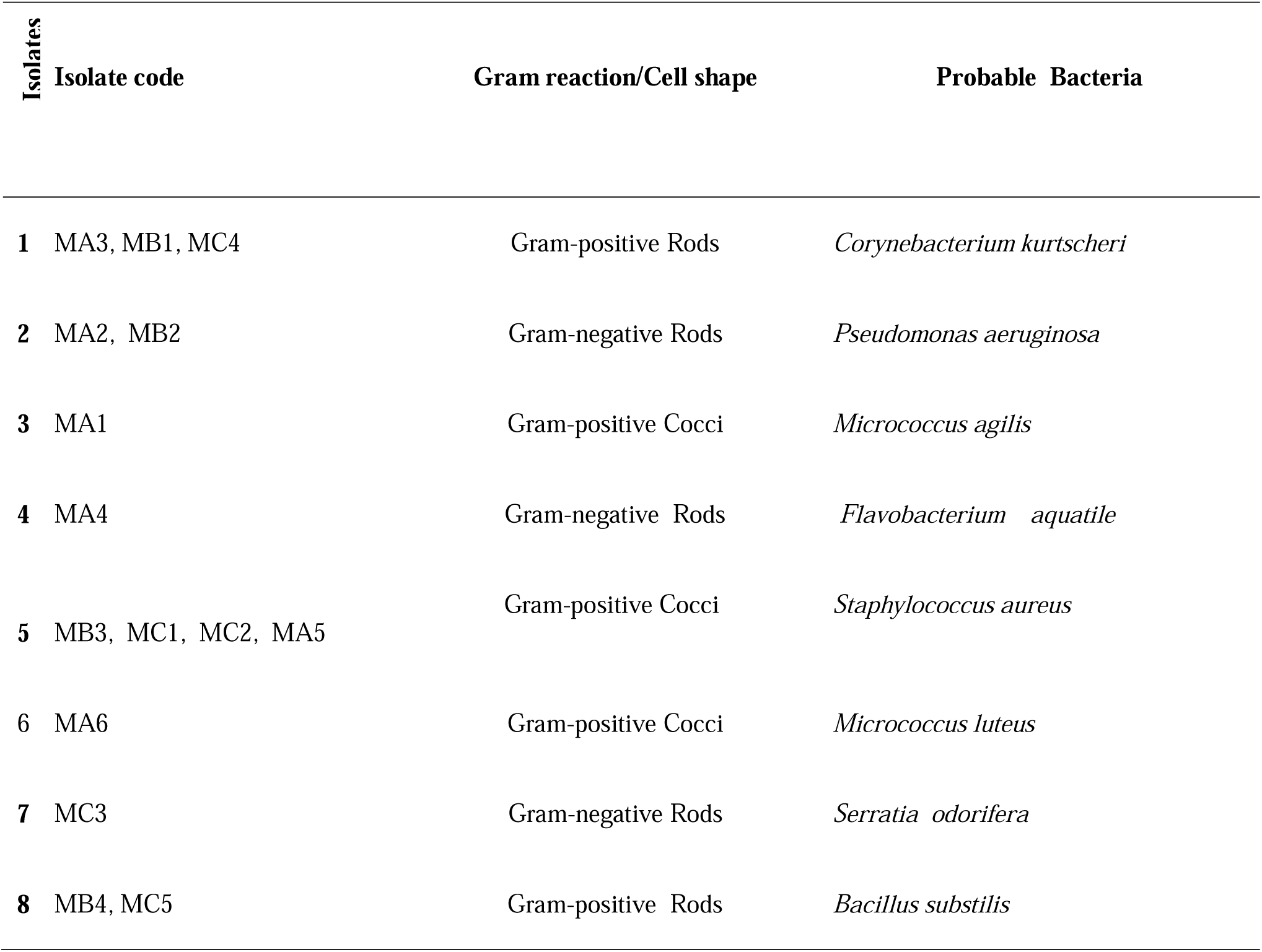
Phenotypic characteristics of bacterial isolates from soil contaminated with waste engine oil.

### Hydrocarbon utilization potential of bacteria

Table 5 shows the changes in pH of MSM during growth of bacteria isolates in hydrocarbon. Decreases in pH (to <pH 7.00) were observed in medium containing of isolates, *Corynebacterium kutscheri, Micrococcus agilis, Serratia odorifera* and *Bacillus substilis*, (which however increased to above pH 7.00). The greatest decrease in pH occurred in cultures of *Corynebacterium kutscheri* (from 7.00 at Day 0 to 5.83 ± 0.34 at Day 15). Medium containing other isolates showed slight increases in pH over time.

**Table 5:**
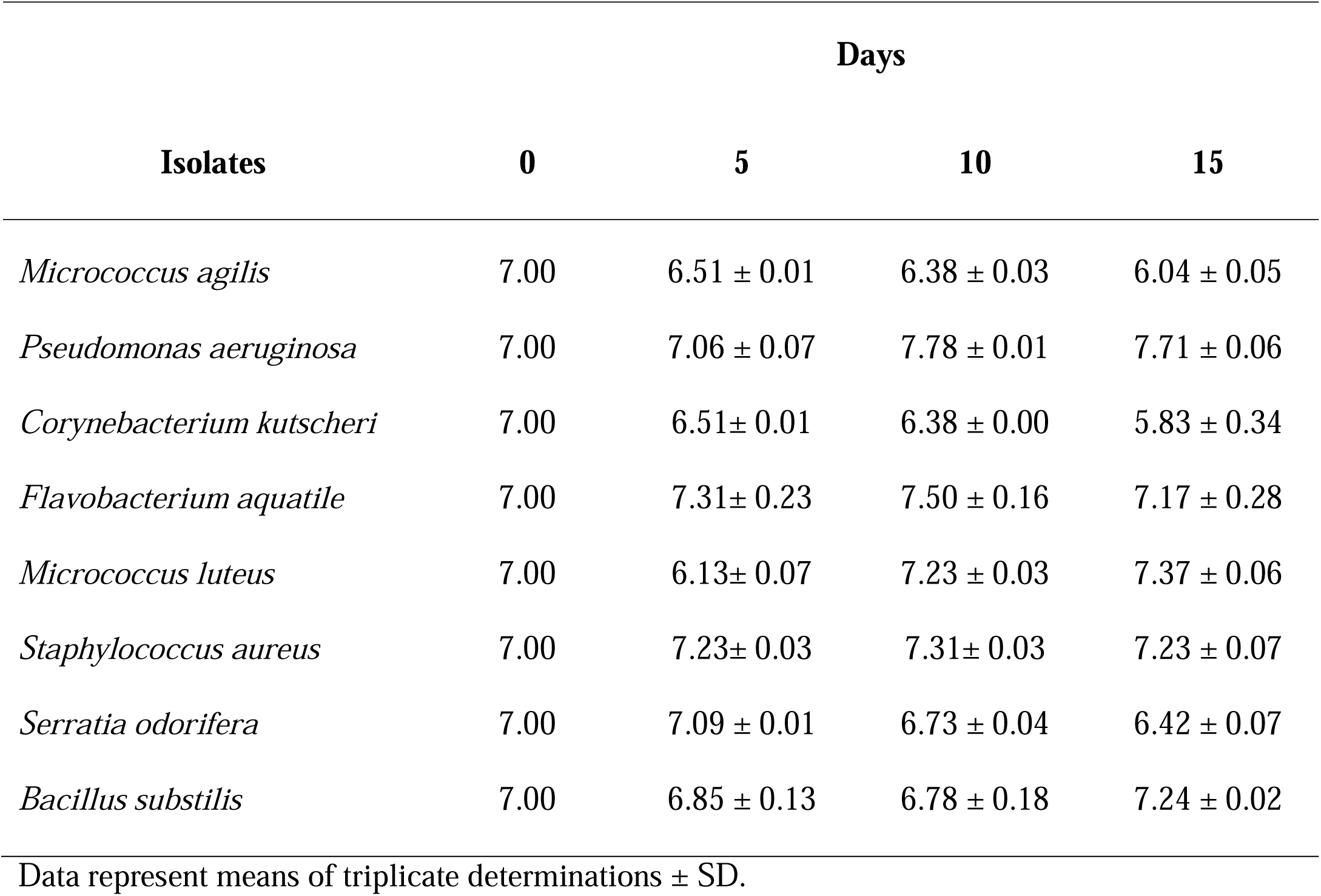
pH of the medium during growth of bacterial isolates in hydrocarbon medium.

| Isolates | Days |  |  |  |
| --- | --- | --- | --- | --- |
|  | 0 | 5 | 10 | 15 |
| *Micrococcus agilis* | 7.00 | 6.51 ± 0.01 | 6.38 ± 0.03 | 6.04 ± 0.05 |
| *Pseudomonas aeruginosa* | 7.00 | 7.06 ± 0.07 | 7.78 ± 0.01 | 7.71 ± 0.06 |
| *Corynebacterium kutscheri* | 7.00 | 6.51± 0.01 | 6.38 ± 0.00 | 5.83 ± 0.34 |
| *Flavobacterium aquatile* | 7.00 | 7.31± 0.23 | 7.50 ± 0.16 | 7.17 ± 0.28 |
| *Micrococcus luteus* | 7.00 | 6.13± 0.07 | 7.23 ± 0.03 | 7.37 ± 0.06 |
| *Staphylococcus aureus* | 7.00 | 7.23± 0.03 | 7.31± 0.03 | 7.23 ± 0.07 |
| *Serratia odorifera* | 7.00 | 7.09 ± 0.01 | 6.73 ± 0.04 | 6.42 ± 0.07 |
| *Bacillus substilis* | 7.00 | 6.85 ± 0.13 | 6.78 ± 0.18 | 7.24 ± 0.02 |

Data represent means of triplicate determinations ± SD.

The changes in the total viable counts of bacterial isolates during 15 days growth in waste engine oil-augmented MSM is shown in Table 6. Total viable counts were higher in cultures containing *Corynebacterium kutscheri* (1.94 ± 0.12 ×10^8^ cfu/ml at Day 5, 6.73 ± 0.45 ×10^8^ cfu/ml at Day 10 and 3.13 ± 0.02 ×10^8^ at Day 15). Growth of *Staphylococcus aureus* showed the lowest decreases (0.31± 0.11 × 10^8^ at Day 5, 0.44 ± 0.01 ×10^8^at Day 10 and 0.42 ± 0.02 ×10^8^at Day 15).

**Table 6:** Total viable count (TVC) of bacterial isolates during growth in hydrocarbon medium (x10^8^ cfu/ml)

| Isolates | Days |  |  |  |
| --- | --- | --- | --- | --- |
|  | 0 | 5 | 10 | 15 |
| *Micrococcus agilis* | 0 | $0.63 \pm 0.03$ | $2.46 \pm 0.02$ | $0.87 \pm 0.06$ |
| *Pseudomonas aeruginosa* | 0 | $0.54 \pm 0.02$ | $2.13 \pm 0.16$ | $0.45 \pm 0.02$ |
| *Corynebacterium kutscheri* | 0 | $1.94 \pm 0.08$ | $5.27 \pm 0.07$ | $3.18 \pm 0.07$ |
| *Micrococcus luteus* | 0 | $0.95 \pm 0.01$ | $2.49 \pm 0.03$ | $1.43 \pm 0.03$ |
| *Flavobacterium aquatile* | 0 | $1.94 \pm 0.12$ | $6.73 \pm 0.45$ | $3.13 \pm 0.02$ |
| *Staphylococcus aureus* | 0 | $0.31 \pm 0.11$ | $0.44 \pm 0.01$ | $0.42 \pm 0.02$ |
| *Serratia odorifera* | 0 | $0.54 \pm 0.03$ | $2.23 \pm 0.25$ | $0.48 \pm 0.04$ |
| *Bacillus subtilis* | 0 | $1.39 \pm 0.41$ | $4.08 \pm 0.65$ | $2.47 \pm 0.07$ |

Data represent means of triplicate determinations ± SD.

Table 7 shows the results of turbidity measurement (OD_600_) of the medium during incubation. The highest increases in turbidity was recorded in cultures with *Corynebacterium kutscheri*, (0.189 ± 0.04 at Day 0 to 0.301± 0.11 at Day 15), and *Bacillus substilis*, (0.165 ± 0.04 at Day 0 to 0.341± 0.02 at Day 15). The lowest turbidity was observed with *Staphylococcus aureus* (0.20 ± 0.04 at day 0 to 0.14 ± 0.02 at day 15).

**Table 7:** Turbidity measurement of bacterial growth in hydrocarbon medium (OD_600_)

| Isolates | Hours |  |  |  |
| --- | --- | --- | --- | --- |
|  | 0 | 5 | 10 | 15 |
| *Micrococcus agilis* | $0.97 \pm 0.01$ | $0.154 \pm 0.01$ | $0.209 \pm 0.12$ | $0.201 \pm 0.03$ |
| *Pseudomonas aeruginosa* | $0.033 \pm 0.001$ | $0.035 \pm 0.10$ | $0.182 \pm 0.10$ | $0.097 \pm 0.05$ |
| *Corynebacterium kutscheri* | $0.189 \pm 0.04$ | $0.252 \pm 0.07$ | $0.386 \pm 0.04$ | $0.301 \pm 0.11$ |
| *Flavobacterium aquatile* | $0.085 \pm 0.01$ | $0.153 \pm 0.08$ | $0.196 \pm 0.07$ | $0.227 \pm 0.05$ |
| *Micrococcus luteus* | $0.040 \pm 0.02$ | $0.083 \pm 0.06$ | $0.162 \pm 0.03$ | $0.183 \pm 0.04$ |
| *Staphylococcus aureus* | $0.20 \pm 0.04$ | $0.053 \pm 0.10$ | $0.084 \pm 0.05$ | $0.014 \pm 0.02$ |
| *Serratia odorifera* | $0.40 \pm 0.02$ | $0.172 \pm 0.11$ | $0.121 \pm 0.001$ | $0.096 \pm 0.01$ |
| *Bacillus subtilis* | $0.165 \pm 0.05$ | $0.204 \pm 0.01$ | $0.413 \pm 0.05$ | $0.341 \pm 0.04$ |

Data represent means of triplicate determinations ± SD.

### Emulsification activity of bacteria

Figure 1 shows the emulsification index (E_24_) of the bacterial isolates in waste engine oil. *Corynebacterium kutscheri* had the highest emulsification index (52 %). The lowest emulsification index was observed for *Staphylococcus aureus* (8 %).

**Figure 1:**
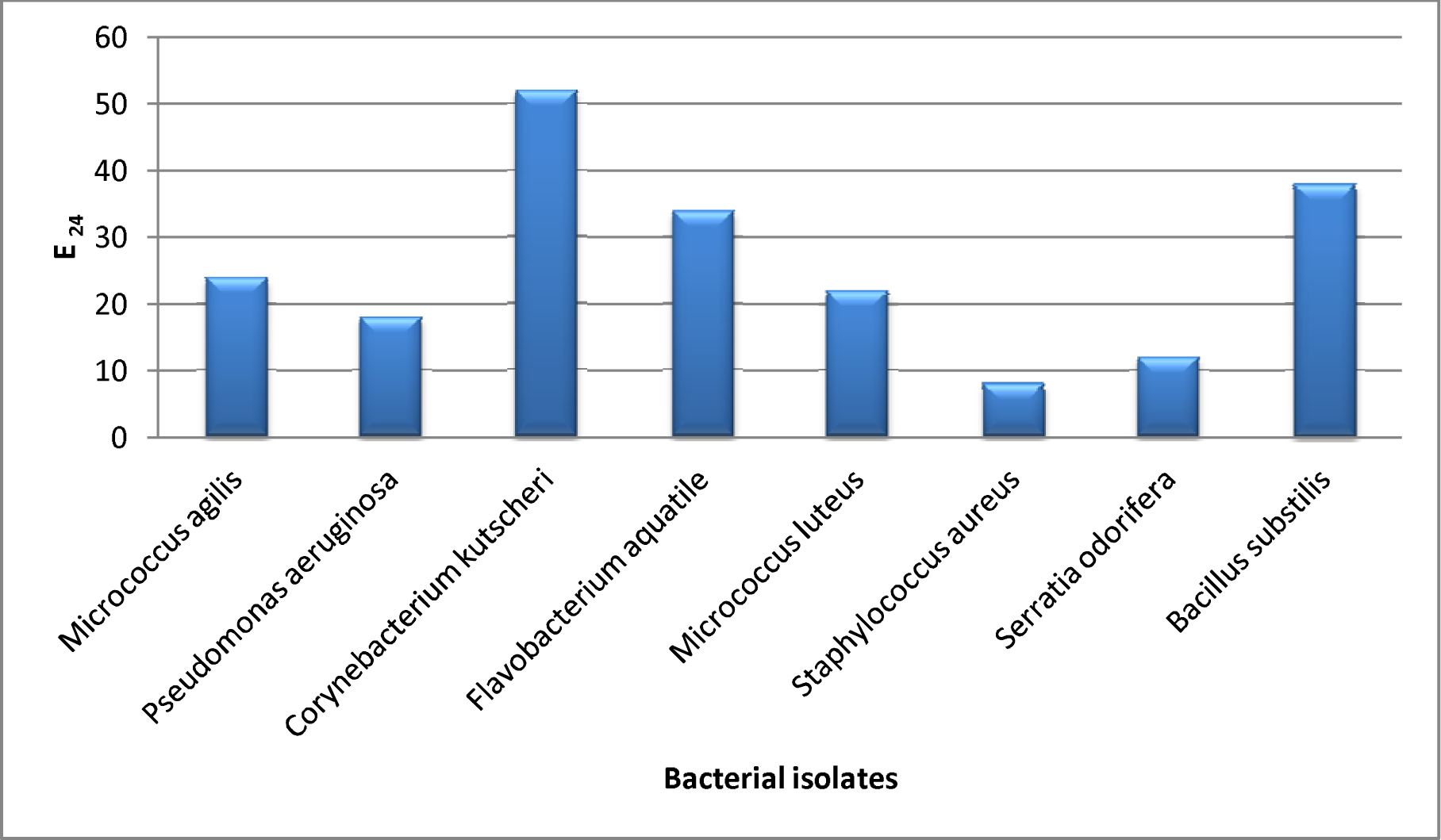
Emulsification activity (E_24_) of bacterial isolates.

## 4. DISCUSSION

This study on the assessment of hydrocarbon utilizing potential of bacteria isolated from waste engine oil-polluted soil on hydrocarbons reveals the presence of hydrocarbon-utilizing bacteria in waste engine oil-polluted soil environment under a variety of experimental conditions..

The results of the physicochemical analysis of the soil samples showed higher levels of properties in the polluted soil samples when compared with the unpolluted soil sample. The results agrees the results of Chikere, [24] and Chikere and Ekwuabu [25] which showed high physicochemical parameters in polluted soil samples compared to unpolluted samples determined and indicated previous exposure of the polluted samples to hydrocarbon contamination with traces of other organic and inorganic contaminants.

The high bacterial counts recorded in polluted soil samples compared with the unpolluted control samples could be attributed to the myriad of nutrients, high organic matter concentration and other ecological factors that influence the survival of soil microorganisms that play important roles in the decomposition and recycling of nutrients [26]. Luepromchai *et al*. [27] has reported an increase in the numbers of hydrocarbon degraders in soil in the presence of PAHs, without any impact on the overall bacterial numbers. The difference between THB and HUB counts was observed to be minimally insignificant which suggests that most of the micro-organisms present in the various polluted sample sites are hydrocarbon degraders that can withstand the concentrations of hydrocarbons and also use them as source of carbon [25].

The result of the emulsification test showed that the isolates produced emulsifying compounds. A large variety and number of biosurfactant producers have been isolated from hydrocarbon-impacted sites [28] although they have also been identified from soils, which are unconnected to hydrocarbon contamination [29]. *Corynebacterium kutscheri* showed the highest emulsification (52 %) at 1% waste engine oil, while the least was *Staphylococcus aureus* (8 %). *Corynebacterium* sp has also been reported by Onuoha *et al.* [30] with high emulsification ability among three hydrocarbon-degrading bacterial species isolated from soil.

## 5. CONCLUSION

This present study illustrates the effects of petroleum hydrocarbon pollution on soil physicochemistry and bacteriology, as well as the potential for using allocthonous bacterial populations in polluted soils for the degradation of petroleum hydrocarbons.

